# Preclinical evaluation of the biodistribution, safety, and hemocompatibility of PEG-coated iron oxide magnetic nanoparticles in a viral oncogenesis mouse model

**DOI:** 10.64898/2026.09.29.755312

**Authors:** Gabriel Principe, Silvina Tiburzi, Virginia Lezcano, Gabriela Montiel Schneider, Flavio Sives, Francisco H Sánchez, Betina N García, Verónica Lassalle, Verónica González-Pardo

**Author notes:** Corresponding author: Verónica González-Pardo, Departamento de Biología, Bioquímica y Farmacia, UNS. San Juan 670, 8000 Bahía Blanca-Argentina. These authors contributed equally (Gabriel Principe, Silvina Tiburzi). **Abbreviations:** ALAT/SGPT, alanine aminotransferase; ALP, alkaline phosphatase; aPTT, activated partial thromboplastin time; ASAT/SGOT, aspartate aminotransferase; H&E, hematoxylin and eosin; HHV-8, human herpesvirus-8; KSHV, Kaposi’s sarcoma-associated herpesvirus; MAG@PEG, polyethylene glycol-coated iron oxide nanoparticles; MNPs, magnetic iron oxide nanoparticles; MPS, mononuclear phagocytic system; MRI, magnetic resonance imaging; PEG, polyethylene glycol; PT, prothrombin time; TT, thrombin time; vGPCR, viral G protein-coupled receptor; VSM, vibrating-sample magnetometer.

## Abstract

Previously, we demonstrated the biosafety of polyethylene glycol-coated iron oxide nanoparticles (MAG@PEG) in a cellular model of viral oncogenesis, supporting their further preclinical evaluation. Herein, we evaluated the safety and biodistribution of MAG@PEG in tumor-bearing mice and assessed the influence of an external magnetic field on nanoparticle distribution. Animals were randomly assigned to three groups: G1, control (PBS); G2, MAG@PEG (10 mg/kg); and G3, MAG@PEG (10 mg/kg) with magnetic exposure for 1 h. Doses were administered intraperitoneally weekly for 1 month. Results showed that animal behavior and body weight were unaffected by MAG@PEG, while tumor growth was significantly reduced only in G2. Biodistribution studies based on tissue magnetization revealed higher magnetic iron levels in the stomach followed by the spleen, lungs, and tumors, particularly in G2. Histological analysis detected Prussian blue-positive staining mainly in the spleen and in lymphoid-associated structures in the lung, stomach and intestine, while a fraction of MAG@PEG remained within the peritoneal cavity. Serum biochemistry showed no evidence of nephrotoxicity. Furthermore, hemocompatibility studies using human blood samples demonstrated that MAG@PEG was non-hemolytic according to the ASTM F756 standard. Taken together, the biodistribution, histological, biochemical, and hemocompatibility findings provide a comprehensive preclinical assessment of the biosafety of MAG@PEG and identify the route of administration and magnetic targeting as key determinants of nanoparticle biodistribution.

## 1. Introduction

The World Health Organization (WHO) and the International Agency for Research on Cancer (IARC) estimate that approximately 10% to 15% of all cancers globally are caused directly by oncoviruses. Seven human oncoviruses have been recognized to develop cancer so far [1]. Viral cancers arise in the context of persistent infections and can manifest years or decades after the acute infection, since these viruses are transmitted between humans and establish chronic, usually asymptomatic infections [2]. Among them, the human herpesvirus-8 (HHV-8) or Kaposi’s sarcoma-associated herpesvirus (KSHV) causes neoplasms characterized by proliferative spindle-cells and angiogenesis [3, 4]. The viral genome encodes the G protein-coupled receptor (vGPCR), which has been demonstrated to drive oncogenesis and angiogenesis [4, 5]. In the last decade, therapeutic choices for Kaposís sarcoma have improved considerably with active antiretroviral therapy, allowing for more effective disease control in numerous cases. However, many patients never complete full remission. Regarding nanomedicine therapies, nearly twenty nanomedicines were approved by the FDA for cancer treatment, and only three were approved for the treatment of Kaposi’s sarcoma: Doxil (Caelyx)®, DaunoXome®, and Lipo-Dox® [6]. Other studies using silver nanoparticles revealed that they cause cytotoxicity in KSHV infected cells, provoking ROS generation and autophagy [7].

In conventional chemotherapy, only a small fraction of the administered drug reaches the tumor, as it is rapidly eliminated from circulation, while causing toxic effects on healthy tissues. In this context, magnetic iron oxide nanoparticles (MNPs) may represent a promising strategy for the treatment of virus-associated cancers. MNPs, especially those based on magnetite and maghemite, have been extensively studied due to their broad potential applications in medicine, particularly in oncology. Their especial properties regarding size, large specific surface area, surface reactivity, and superparamagnetism enable their use as nanotheranostic for targeted drug delivery and as contrast agent in magnetic resonance imaging (MRI) [8]. The exceptionality of MNPs lies in their superparamagnetic nature. When exposed to an external magnetic field, they become strongly magnetized and behave as small independent magnets. The external magnetic field can be applied as stimuli allowing the guidance of MNPs to specific locations, enabling active targeted drug delivery and potentially reducing both systemic toxicity and required therapeutic dose. Superparamagnetism distinguishes MNPs from other types of nanoparticles used in medicine, such as gold, silver, or polymer nanoparticles. In this regard, metallic nanoparticles have been proven to be non-biodegradable and toxic. In addition, the toxicological profile and active targeting capabilities of most polymer nanoparticles have not yet been thoroughly evaluated [9]. In contrast, MNPs are mainly biodegradable. Evidence indicates that MNP biodegradation involves lysosomal enzymes that release free iron ions. Subsequently, these ions can be stored as ferritin and hemosiderin for future cell use. MNP biocompatibility is attributed to an effective iron metabolism allowing iron recycling and excessive iron exocytosis. Many studies have shown that MNPs present low toxicity; nevertheless, their cytotoxicity largely depends on their size, shape, surface coating, concentration, exposure time, and cell type which they interact to [10]. In line with these findings, our group recently demonstrated that MNPs coated with polyethylene glycol (PEG) at concentrations above 50 μg/mL of MNPs increased metabolic and lysosomal activity as well as ROS production, without affecting vGPCR cell proliferation under the tested conditions [11]. Previous *in vivo* studies have also reported low toxicity associated with MNP exposure, with only transient effects such as oxidative stress. Pathological alterations in the liver and spleen have been reported only at extreme doses (500 mg Fe/kg) [9]. Collectively, these findings suggest that MNPs may induce transient oxidative stress without causing permanent cellular or organ damage, depending on the administered dose.

MNP Biodistribution is strongly influenced by their physicochemical properties, including particle size, surface coating, and surface charge. In addition, the route of administration plays a critical role in determining their distribution and clearance. MNPs are commonly administered intravenously when used as MRI contrast agents or for oncological application [8, 12].

Once bare MNPs enter the bloodstream, proteins, complement components, and other serum biomolecules rapidly adsorb onto their surface as result of hydrophobic and electrostatic interactions, forming a protein corona [13]. This process largely determines their biological fate, including their recognition and uptake by macrophages of the mononuclear phagocytic system (MPS), formerly known as reticuloendothelial system, as well as their renal clearance. In addition, bare MNPs are preferentially accumulated in the liver and spleen, two major organs of the MPS involved in MNP sequestration and elimination. At high intravenous doses, the excess of MNPs may also accumulate in other tissues, including the lungs, heart, and adipose tissue [9]. Functionalization with bioactive compounds and biocompatible coating improves nanoparticle stability and reduces aggregation. In this regard, PEG is widely recognized as an effective coating material because of its excellent biocompatibility and its ability to reduce phagocytic uptake, thereby prolonging nanoparticle circulation time [14, 15].

Hemocompatibility is a critical consideration in the design of new MNP-based systems to facilitate their successful translation into clinical applications. In this regard, Malehmir and collaborators, highlighted that the surface chemistry of MNP is a key determinant of their hemocompatibility. Specifically, the interaction of MNPs with blood components and their overall hemocompatibility profile depend on several factors, including surface charge, geometry, porosity, and the surface functionalization type, such as polymers coating or specific functional groups [16].

In our previous study, we investigated the biological effects of PEG-coated iron oxide magnetic nanoparticles (MAG@PEG) in a cellular model of Kaposi’s sarcoma, in which endothelial cells stably expressing the vGPCR oncogene undergo malignant transformation. Furthermore, the physicochemical characterization and biological evaluation of MAG@PEG demonstrated favorable properties for the biomedical applications.

As a further step toward preclinical development, the present study aimed to evaluate the biodistribution, tumor-targeting capacity, biosafety, and hemocompatibility of MAG@PEG in a murine model of Kaposís sarcoma.

## 2. Materials and Methods

### 2.1. Chemicals and Reagents

All reagents were of analytical grade and used without further purification. Ferric chloride hexahydrate was purchased from Tetrahedron (Mendoza, Argentina). Ferrous sulphate heptahydrate was provided by Mallinckradt Chemical Works (t. Louis, MO, USA). Polyethylene glycol (PEG 6000) was acquired from Biopack (Zárate, Argentina). Sodium hydroxide was provided by Cicarelli (Santa Fe, Argentina). Hematoxylin and eosin (H&E) stains were gently donated by Dr. María Constanza Díaz Andrade (Lab. Citología, Histología y Embriología Animal, INBIOSUR, CONICET-UNS). Commercial diagnostic kits for prothrombin time (Soluplastin), fibrinogen, activated partial thromboplastin time (APTTest) and thrombin time were gently donated by Wiener lab. (Rosario, Santa Fe, Argentina).

### 2.2. MAG@PEG synthesis and preparation for biological assays

The MAG@PEG nanosystem was synthesized following previously optimized procedures using the hydrothermal method [11, 17]. Briefly, PEG 6000 (0.5965 g) was dissolved in 30 mL of distilled water. Subsequently, iron salts (0.6687 g of Fe^3+^ salt and 0.3607 g of Fe^2+^ salt) were added to the solution and stirred for 10 min at room temperature. Then, a 5 M NaOH solution was added until the pH exceeded 10. The reaction mixture was transferred to a Teflon-lined autoclave and maintained at 160 °C for 9 h, followed by cooling at room temperature. Afterwards, the nanosystem was decanted using a neodymium magnet, and the supernatant was removed. Finally, MAG@PEG were washed with distilled water until the pH and conductivity values were similar to those of distilled water.

A MAG@PEG stock suspension (1 mg/mL) was prepared in physiological saline (0.9% NaCl) and dispersed by sonication cycles immediately before use. The MAG@PEG concentration used in the *in vitro* hemocompatibility assays was estimated from the nanoparticle dose administered *in vivo* (10 mg/kg, groups G2 and G3). To approximate the corresponding blood concentration, the total amount of MAG@PEG administered per mouse was calculated assuming instantaneous distribution throughout the average mouse blood volume (approximately 1.5 mL). Based on this estimation, a MAG@PEG concentration of 160 µg/mL was selected for *in vitro* testing. Additional concentrations of 16 and 320 µg/mL were also included to evaluate hemocompatibility over a broader concentration range.

### 2.3. Animals

Six-week-old N:NIH(S)-nu mice (17-21 g) were acquired from *Laboratorio de Animales de Experimentación* (La Plata, Argentina) and housed in the animal care facility of the Department of Biology, Biochemistry, and Pharmacy of the National University of the South (DBByF-UNS) in accordance with the National Research Council’s Guide for the Care and Use of Laboratory Animals [18]. The animals were kept in pathogen free cages under constant room temperature (22 ± 1 °C) and humidity (70%), with a 12 h light/dark cycle, and provided with sterile water and a standard autoclaved diet *ad libitum*. The employed protocol was approved and registered under protocol N° CDBByF 701/23 by the Institutional Committee for the Care and Use of Experimental Animals of DBByF-UNS.

### 2.4. Tumor allografts and experimental design

Tumor allografts were generated as previously described by our group [19]. Briefly, vGPCR cells (2 x 10^6^ in 0.2 mL DMEM with 5% FBS and 500 µg/mL G418) were injected subcutaneously into the right flank of female nude mice. Tumor volume was estimated measuring tumor length and width with a caliper to apply the formula: tumor volume (mm^3^) = [(width)^2^ x length]/2. When the tumors had reached a volume of approximately 100 mm^3^, mice were divided into three groups of four animals each (n = 4). The treatment was administered by intraperitoneal (IP) injection once a week for one month; the control group (G1) received the vehicle solution (PBS), whereas the treatment groups (G2 and G3) received MAG@PEG at a dose of 10 mg/kg. Additionally, a 0.3 Tesla neodymium magnet was exposed over the skin in the tumor proximity for 1 h in the mice corresponding to group 3 (G3). Mice weight and behavior were monitored throughout the experience as a parameter of toxicity. At the end of the experiment, animals were anesthetized with Ketamine/Xylazine (80 mg/kg + 10 mg/kg) to collect the blood and thereafter euthanized by decapitation. Then, the tumors and different organs (spleen, lungs, heart, stomach, kidneys, liver, intestine, and brain) were harvested. In addition, tumors were weighted and photographed.

### 2.5. Animal blood collection and biochemical parameters analysis

Blood samples were collected from sedated mice via intracardiac puncture using a 1 mL syringe fitted with a 25G needle. The blood (500 - 800 µL) was collected in Eppendorf tubes and sent for analysis to *Bioquímica Austral, Laboratorio de Análisis Clínicos y Gestión* (Bahía Blanca, Buenos Aires, Argentina). Glucose, urea, creatinine, alkaline phosphatase (ALP), alanine aminotransferase (ALAT/SGPT), and aspartate aminotransferase (ASAT/SGOT) were analyzed. The biochemical kits and calibration controls were from Wiener lab. (Rosario, Santa Fe, Argentina) and used according to manufacturer’s protocols.

### 2.6. Quantification of MAG@PEG by VSM

To quantify the MAG@PEG content in the selected tissues, the isothermal magnetization was measured using a LakeShore 7404 VSM. Measurements were performed *ex vivo* on dehydrated tissues that were stored frozen and exposed to the ambient environment only during handling. Transport and experimental measurements were conducted at room temperature (approximately 15 - 22 °C), with the material carefully preserved using appropriate wrapping. Special care was taken to avoid contamination, particularly with magnetic substances. When tissues exceeded the magnetometer capacity, they were sectioned when possible and measured separately. The overall result corresponding to the complete tissue was subsequently calculated. Otherwise, the tissues were homogenized in an agate mortar and pestle and a representative aliquot was analyzed.

### 2.7. Histological analysis

Tissues were excised and fixed in 10% neutral buffered formalin for 24 h at room temperature. Subsequently, samples were dehydrated through a graded ethanol series, and cleared in n-butanol prior to paraffin embedding. Paraffin-embedded tissues were sectioned using a microtome at 5 µm thickness. Sections were stained with hematoxylin and eosin (H&E) and Prussian blue, then examined under a bright-field microscope [20]. The same procedure was applied to samples obtained from the clot formation assay after exposure to MAG@PEG, prior to histological evaluation.

### 2.8. Human blood collection

Peripheral venous blood samples were collected from healthy adult volunteers into EDTA or sodium citrate anticoagulant tubes or without anticoagulant tubes after obtaining written informed consent. All procedures involving human samples were approved by the Research Ethics Committee of the Hospital Italiano Regional del Sur (Bahía Blanca, Argentina), registered in the Provincial Registry of Research Ethics Committees of the Ministry of Health of the Province of Buenos Aires (approval N° 020/2010). Samples were processed within 2 h after collection. Coagulation parameters were determined using commercial diagnostic kits for prothrombin time (Soluplastin), fibrinogen, activated partial thromboplastin time (aPTTest) and thrombin time according to the manufacturer’s instructions.

### 2.9. Hemocompatibility assays

#### 2.9.1. Hemolysis assay

MAG@PEG hemolytic activity was evaluated using EDTA-anticoagulated whole blood. Blood samples were incubated with physiological saline (negative control), distilled water (positive control), or different concentrations of MAG@PEG (16 µg/mL, 160 µg/mL, and 320 µg/mL). The concentrations of MAG@PEG used were selected to approximate doses of 1, 10, and 20 mg/kg, respectively. Samples were incubated for 1 h at 37 °C in a shaking water bath. Thereafter, samples were centrifuged at 1,500 rpm for 10 min to obtain plasma. The plasma containing released hemoglobin was collected and aliquoted (100 µL) in triplicate into a 96-well microplate and absorbance was measured at 540 nm with a microplate reader. Hemolysis was expressed as a percentage of hemoglobin release relative to negative and positive controls according to the ASTM F756-17 standard, using the following equation:

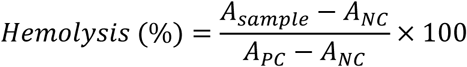

where *A_sample_*, *A_NC_*, and *A_PC_* correspond to the absorbance of the sample, negative control, and positive control, respectively.

#### 2.9.2. Coagulation assays

MAG@PEG effect on blood coagulation was evaluated using citrated whole blood. Blood samples were incubated with physiological saline solution (control), or different concentrations of MAG@PEG (16 µg/mL, 160 µg/mL and 320 µg/mL) for 1 h at 37 °C in an incubator shaker. After incubation, samples were centrifuged at 1,500 rpm for 10 min to obtain plasma. Coagulation parameters were determined using commercial diagnostic kits (Wiener lab, Rosario, Argentina) according to the manufacturer’s instructions.

### 2.10. Statistical analysis

Data normality was evaluated using the Shapiro–Wilk test, while homoscedasticity was assessed through the Levene test. Multiple comparisons were made using one-way ANOVA followed by Dunnett’s test or the non-parametric Mann–Whitney U test. Linear mixed-effects models were used to evaluate tumor growth and mice weight progression throughout the experiment. Statistical significance is shown as *p < 0.05; **p < 0.01.

## 3. Results

### 3.1. Effect of MA@PEG on tumor development

Magnetic nanotheranostics represent a promising alternative to conventional approaches for several pathologies. As mentioned in the Introduction, our previous study demonstrated that the MAG@PEG nanosystem is an efficient and biocompatible drug carrier with magnetic targeting capabilities that could be used to reduce adverse effects on healthy tissues [11]. The physicochemical, structural and stability properties of this nanosystem have been extensively characterized, supporting its further evaluation *in vivo* assays [17].

To further explore these findings and assess the potential of this nanosystem as a nanotheranostic platform, an experimental study was designed to evaluate the biodistribution, biosafety, and magnetic targeting of intraperitoneally administered MAG@PEG in an animal model of Kaposi’s sarcoma (**Figure 1**). Briefly, the vGPCR cells were subcutaneously injected in the right flank of female nude mice, which were randomly divided in three groups of four animals each when tumors reach a volume of approximately 100 mm^3^. The dose and route of administration of MAG@PEG were chosen based on the preclinical evaluation of a similar nanosystem, composed of chitosan-coated iron oxide magnetic nanoparticles (MNPs) [21]. As shown in **Figure 1**, mice were treated by weekly intraperitoneal injection for 30 days with PBS (G1: Control), a suspension of 10 mg/kg of MAG@PEG (G2: MAG@PEG), or the same treatment as G2 including a 0.3 Tesla magnet exposed externally over the skin at the tumor site for 1 h (G3: MAG@PEG + Magnet). During the experiment, animal behavior and weight were monitored as indicators of toxicity, and neither of these parameters were altered during the test (**Figure 2A**). In **Figure 2B**, representative photographs of tumor progression in one mouse from each group throughout the experimental period are shown (selected from **Suppl. Fig. 1**). Additionally, tumor volume was measured during the study. In **Figure 2C**, G2 exhibited delayed tumor development compared to the control (G1) and the magnet-exposed group (G3), with highly significant differences versus G1 (p < 0.01) and significant differences versus G3 (p < 0.05). Unexpectedly, no significant differences were observed between G1 and G3. At the end of the experiment, animals were anesthetized for blood collection and subsequently euthanized. Tumors and selected organs were then collected. As shown in **Figure 2D**, tumor weight tended to decrease in G2 and G3; however, these differences were not statistically significant compared to G1.

**Figure 1.**
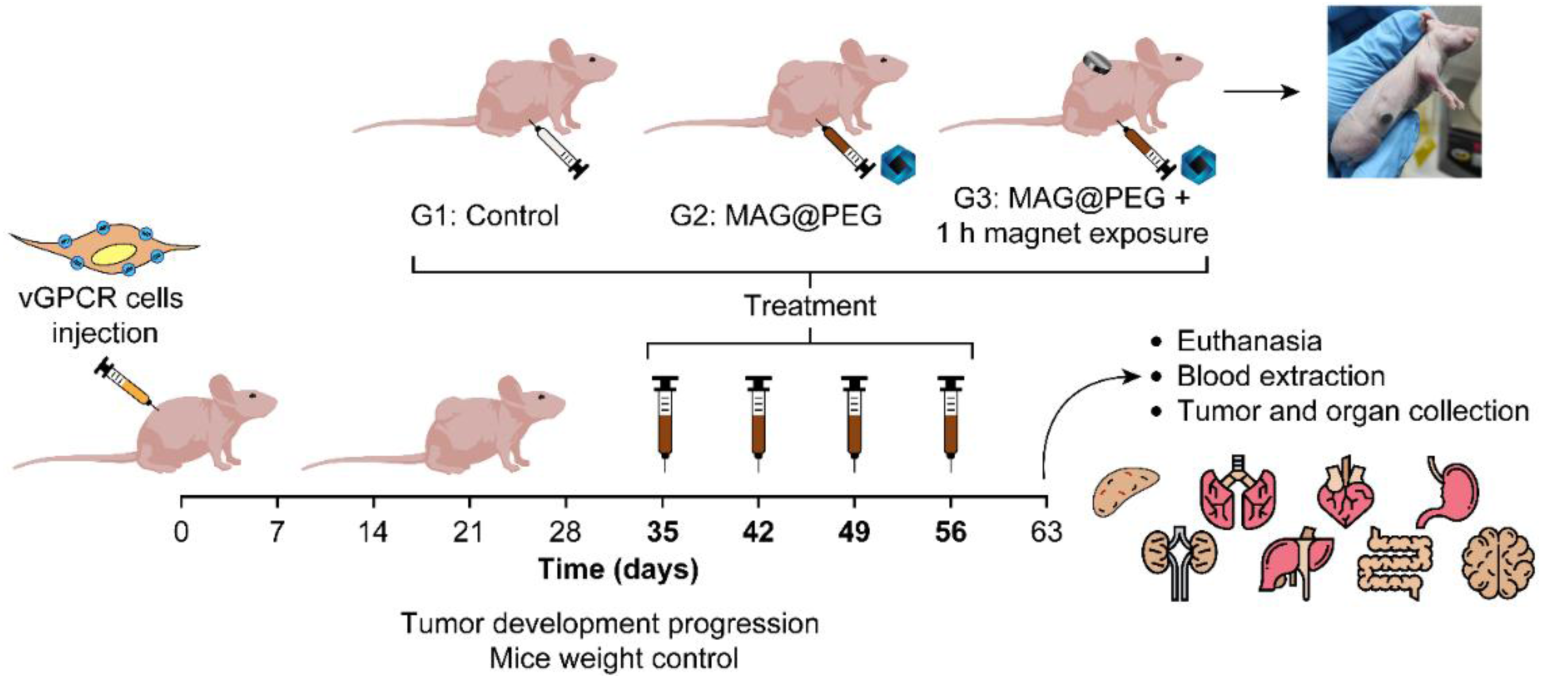
Tumor allografts and experimental design. Female nude mice were injected subcutaneously with vGPCR cells to generate the animal model of Kaposi’s sarcoma. When the tumors reached a volume of 100 mm^3^, mice were divided into three groups and received PBS (G1), MAG@PEG (10 mg/kg) (G2), and MAG@PEG (10 mg/kg) plus the attachment of a 0.3 T magnet over the tumor for 1 h (G3). The doses were administered via intraperitoneal injection once a week for one month. At the end of the experiment, mice were euthanized, blood samples were collected, and tumors and different organs were harvested.

**Figure 2.**
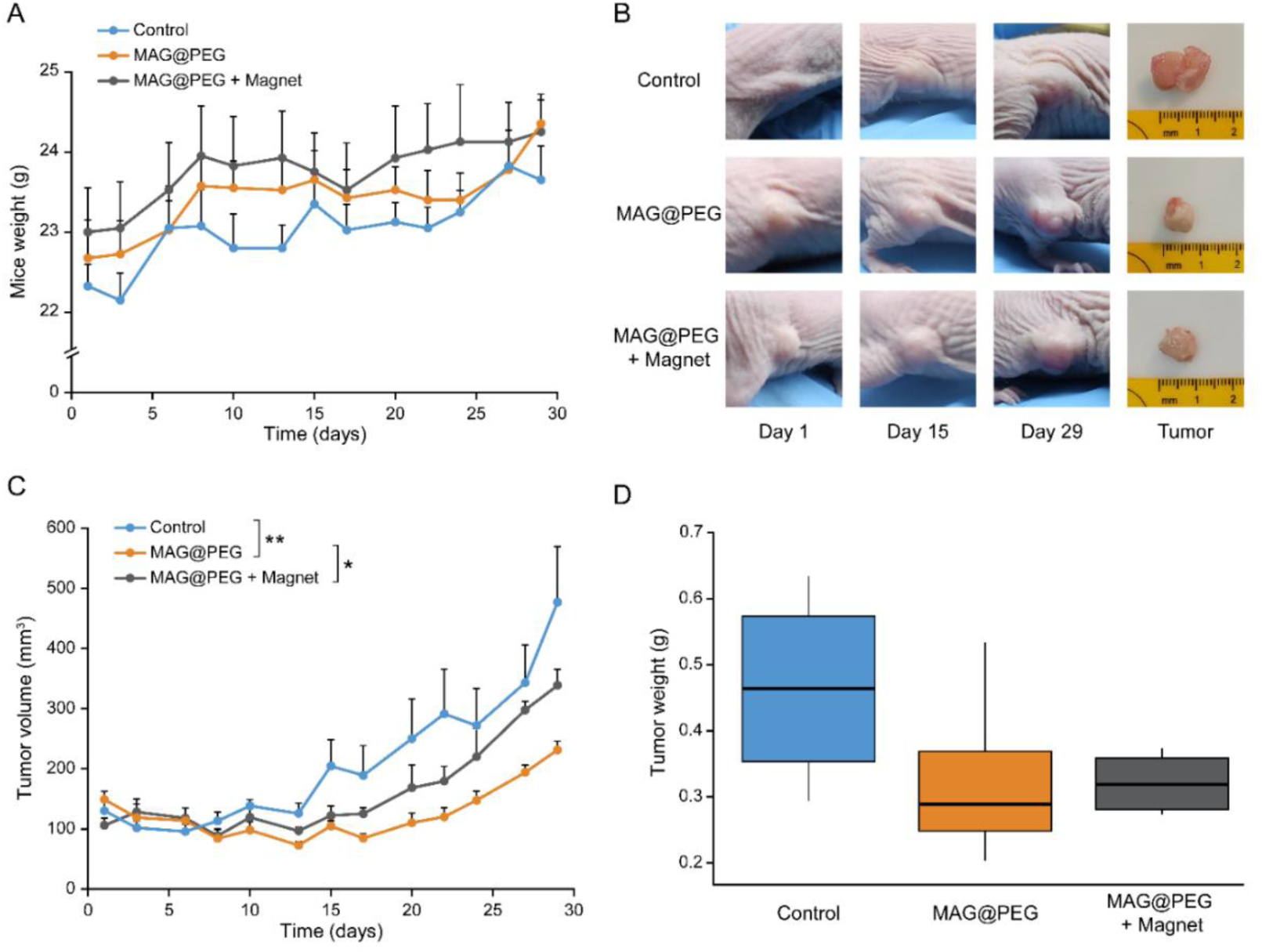
Tumor development analysis. (A) Mice’s weight progress from the beginning of the treatment until the end of the experiment. (B) Representative photographs of tumor progression in one mouse of each group throughout the experimental period (day 1, 15 and 29) and a photograph of the extracted tumors (Tumor). (C) Tumor growth graph from the beginning until the end of the treatment. (D) Extracted tumor weight. (A) and (C) Data are presented as means ± SEM, statistically significant differences were evaluated using a linear mixed effects model (*p < 0.05; **p < 0.01). (D) No statistically significant differences were found using Dunnett’s test.

The serum samples were analyzed to assess hepatic and renal function (**Table 1**). No statistically significant differences were observed in glucose, urea, creatinine, or ALP among the groups. However, a significant increase in liver transaminases (ALAT and ASAT) was detected in G3 suggesting potential hepatocellular damage. Regarding renal function, creatinine levels remain unchanged across groups, while a slight, non-significant increase in urea was observed in G3, which may reflect mild functional alterations without evidence of structural renal damage (**Figure 5**). In addition, blood smears analysis revealed no alterations in erythrocyte morphology in any group (**Suppl. Fig. 2**).

**Table 1.** Effect of MAG@PEG treatment on serum biochemical parameters in mice. Serum samples were collected for the determination of glucose, urea, creatinine, alkaline phosphatase (ALP), alanine aminotransferase (ALT/SGPT), and aspartate aminotransferase (AST/SGOT) levels. Data are presented as mean ± SEM. Statistical significance was assessed by one-way ANOVA followed by Dunnett’s *post hoc* test for multiple comparisons against the control group (*p < 0.05).

| Parameter | G1: Control | G2: MAG@PEG | G3: MAG@PEG + Magnet |
| --- | --- | --- | --- |
| Glucose (mg/dL) | 277 ± 52.6 | 322.5 ± 57.3 | 272.5 ± 13.3 |
| Urea (mg/dL) | 56.5 ± 6.7 | 56.5 ± 5.4 | 65.5 ± 6.7 |
| Creatinine (mg/dL) | 0.33 ± 0.05 | 0.4 ± 0.04 | 0.34 ± 0.04 |
| ALP (U/L) | 176 ± 49.1 | 206.2 ± 29.3 | 188.5 ± 15.9 |
| ALAT/SGPT (U/L) | 32.8 ± 4.1 | 40.2 ± 7.8 | **59.25 ± 25.3 \*** |
| ASAT/SGOT (U/L) | 76.5 ± 4.5 | 81.25 ± 3.8 | **111 ± 33.9 \*** |

### 3.2. Biodistribution and accumulation of MAG@PEG

To determine the biodistribution and accumulation of MNPs in tumors and organs, magnetization analysis was performed using a VSM with a sensitivity of 5 x 10⁻⁶ emu, implying that amounts greater than 100 ng of MAG@PEG can be reliably detected and quantified (**Figure 3**). Considering the sample size and the MNP saturation magnetization, this corresponds to a minimum detectable MNP mass fraction of about 10⁻⁶ g/g. Because some tissues were expected to contain very low MNP mass fractions, simple fitting models were preferred over more complex alternatives. Instead, an analysis of the asymptotic behavior of the magnetization (M) vs. applied field (H) curves was adopted. Taking advantage of the fact that the experimental curves represent a superposition of the MNP signal (essentially superparamagnetic) and the tissue signal (diamagnetic), the high-field regions were fitted with linear functions, requiring the slopes to be equal in the positive and negative field zones. The saturation magnetization (Mₛ) was estimated as half the difference between the y-intercepts of the two fitted lines. The MNP mass fraction was obtained by calculating the ratio between the Mₛ of the tissue sample and the Mₛ of a sample composed exclusively of MNP powder, as

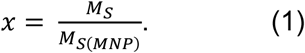

**Figure 3.**
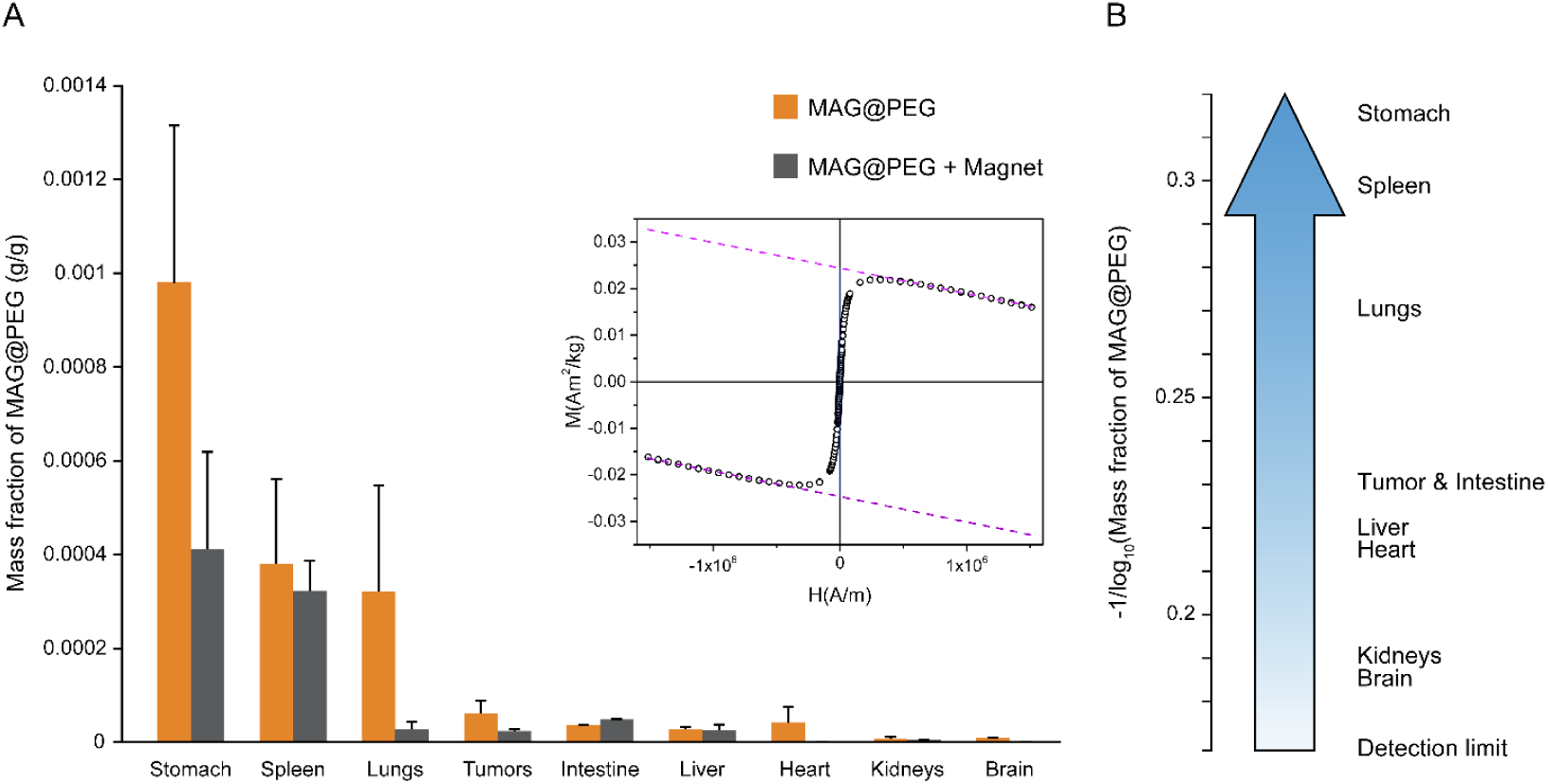
MAG@PEG biodistribution analysis. (A) The graph displays MAG@PEG’s mass fraction in each organ/tumor as mean ± SEM, and an insert representative graph of M vs. H curves of a mouse spleen of G3. The control group is not shown as the mass fraction is virtually negligible. (B) The graph shows the differential uptake of MAG@PEG across the analyzed organs on a logarithmic scale regardless of the application of the magnet, including the detection limit of the technique.

The asymptotic, high-field regions were extensive because the MNPs saturate at a low field compared to the maximum value of 19 kOe (see inset in **Figure 3**). This results in a reliable linear fit and reduces the error arising from the statistical dispersion of the experimental data points.

The global magnetization M_G_ of a subset of samples was defined as the mass-weighted average of the individual values M_i_,

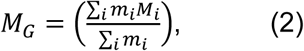

where *m_i_* is the mass and *M_i_* the magnetization of a tissue sample from a mouse belonging to a given group. As a weighted mean value, *M_G_* has a standard dispersion *σ_MG_* given by

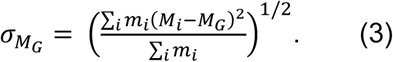

The bars and vertical lines in **Figure 3A** represent the values obtained using equations (2) and (3), respectively. In each case, the results were presented in the following order: MAG@PEG (G2), and magnet-exposed group (G3), as explained in the experimental design (**Figure 1**). The control group was not included as the signals from all tissues were virtually negligible. In contrast, **Figure 3B** presents the differential uptake of MNPs across the analyzed tissues for all mice that received MNPs (G2 and G3 combined) on a logarithmic scale. As shown in **Figure 3**, MAG@PEG accumulation was highest in the stomach, spleen, and lungs, followed by tumors and other organs. Additionally, as can be seen in **Figure 3A**, magnetization measurements were generally higher in G2 than in G3, even in the tumor measurements.

Eq. (1) relies on the assumption that MNPs retain their magnetic properties, in particular, the saturation magnetization M_S,_ after injection, distribution, and relocation into the various tissues. To examine this hypothesis, two additional determinations were performed. Firstly, a second estimation of the mass fraction using *x′* = *χ*/*χ*(MNP), where *χ* is the susceptibility of the superparamagnetic contribution in the quasi-linear low-field region was calculated (see **Suppl. Material**, **Suppl. Fig. 3.I**). It was found that the correlation between *x′* and x is strongly linear, with a slope of approximately 0.95 (see **Suppl. Material**, **Suppl. Fig. 3.II**). This linear correlation between M_S_ / M_S_(MNP) and *χ*/*χ*(MNP), with a slope close to unity, provides strong evidence that the MAG MNPs retain their magnetic properties. Secondly, for tissues exhibiting a sufficiently intense magnetic signal from the MNPs, coercivity values were determined and compared with those observed for the MAG powder (see **Suppl. Material**, **Suppl. Table 1**). The average coercivity across tissues was 2595 A/m, with a standard deviation of 206 A/m. The percentage difference between this average and the coercivity of the MNP powder (2664 A/m) was less than 3%. Together, these two analyses confirm that the MAG MNPs retain their magnetic properties to a sufficiently acceptable degree for a reliable estimation of their mass fraction in tissues via Eq. (1) (see **Suppl. Material**, **Suppl. Fig. 3.III**).

Furthermore, histological analysis of tumors and organs of interest was performed to evaluate the distribution of MAG@PEG and potential tissue damage induced by the nanosystem (**Figures 4** and **5**). As shown in **Figure 4**, tumor sections stained with H&E and Prussian blue, which detects iron-containing deposits, exhibited mainly preserved histological architecture, with Prussian blue-positive stains in G3. Likewise, no major histopathological alterations were observed among the tissue sections of the analyzed organs (**Figure 5**). Prussian blue-positive staining was consistently found only in the spleen across all groups; however, quantitative analysis revealed a higher stained area in G2 and G3 compared to G1 (**Suppl. Fig. 4**). Interestingly, additional Prussian blue-positive deposits were observed in lymphoid-associated tissues, including liver, lymphoid follicles of lungs, and intestine in G2 as well as lymphoid-adipose structures associated with the stomach in G3 (**Figure 6**).

**Figure 4.**
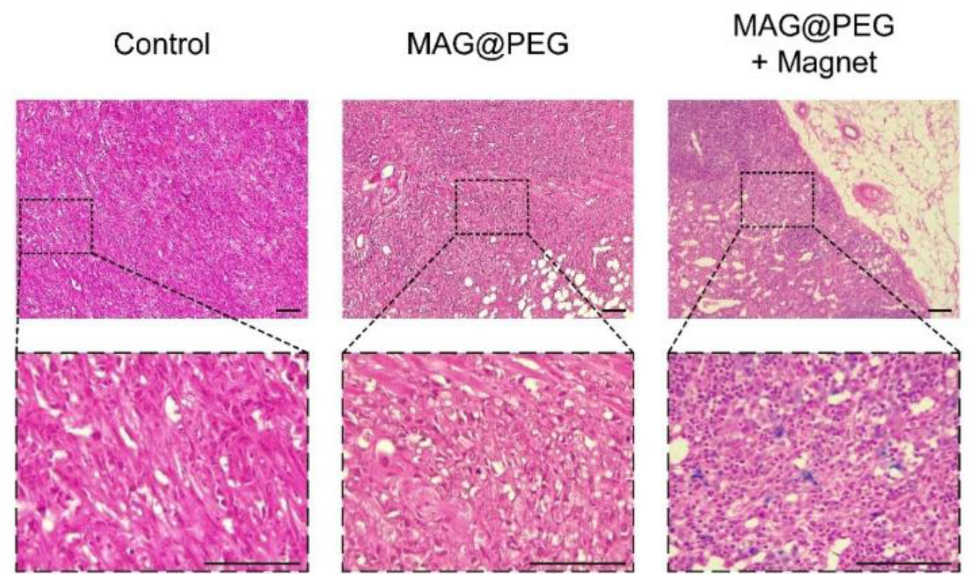
MAG@PEG detection in tumor sections. Micrographs of tumor sections stained with H&E and Prussian blue. Small Prussian blue-positive clusters were only found in G3 tumors. Scale bar: 100 µm.

**Figure 5.**
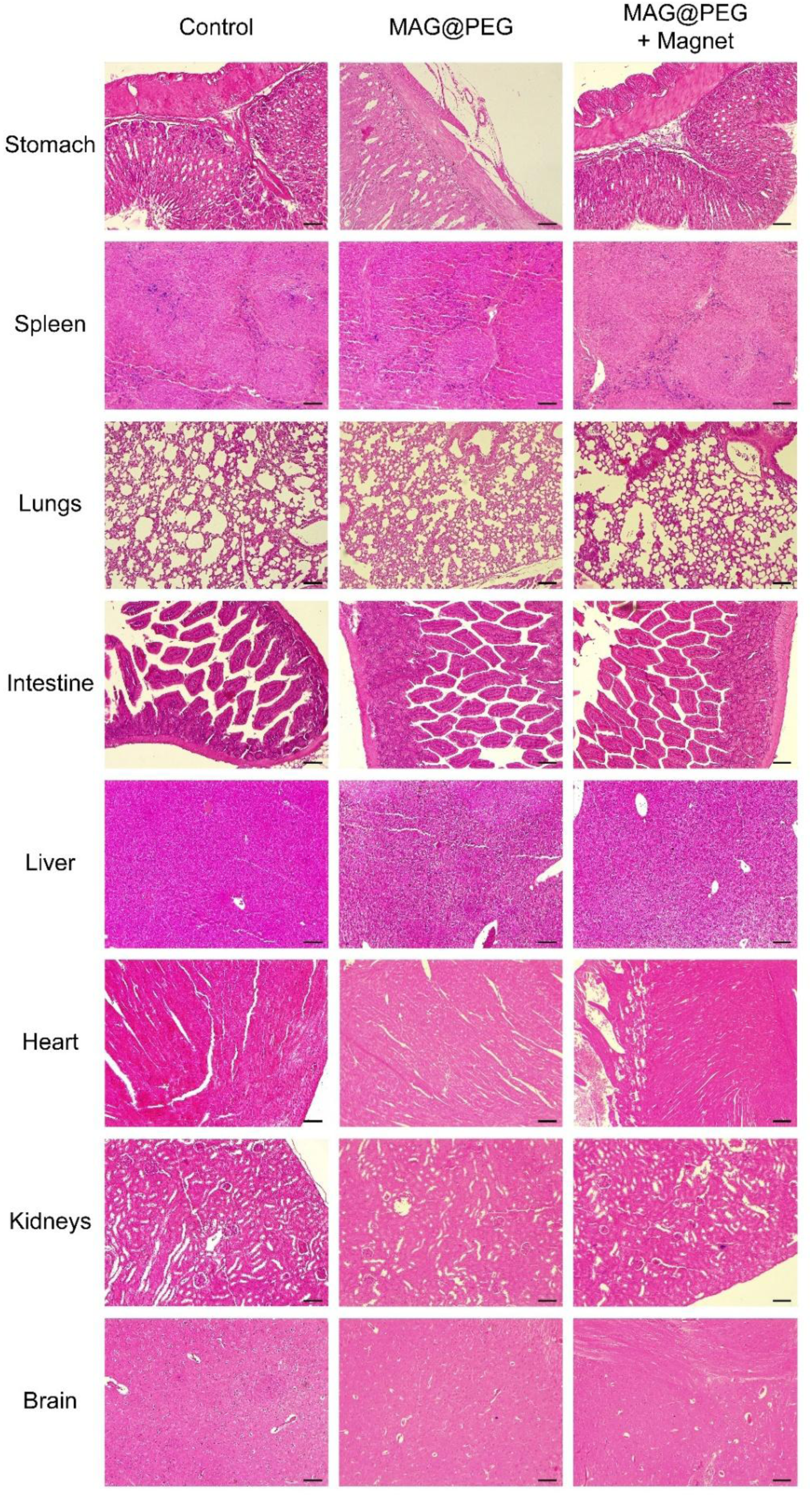
MAG@PEG biodistribution analysis on tissue organs. Representative micrographs of the extracted organs stained with H&E and Prussian blue. Scale bar: 100 µm.

**Figure 6.**
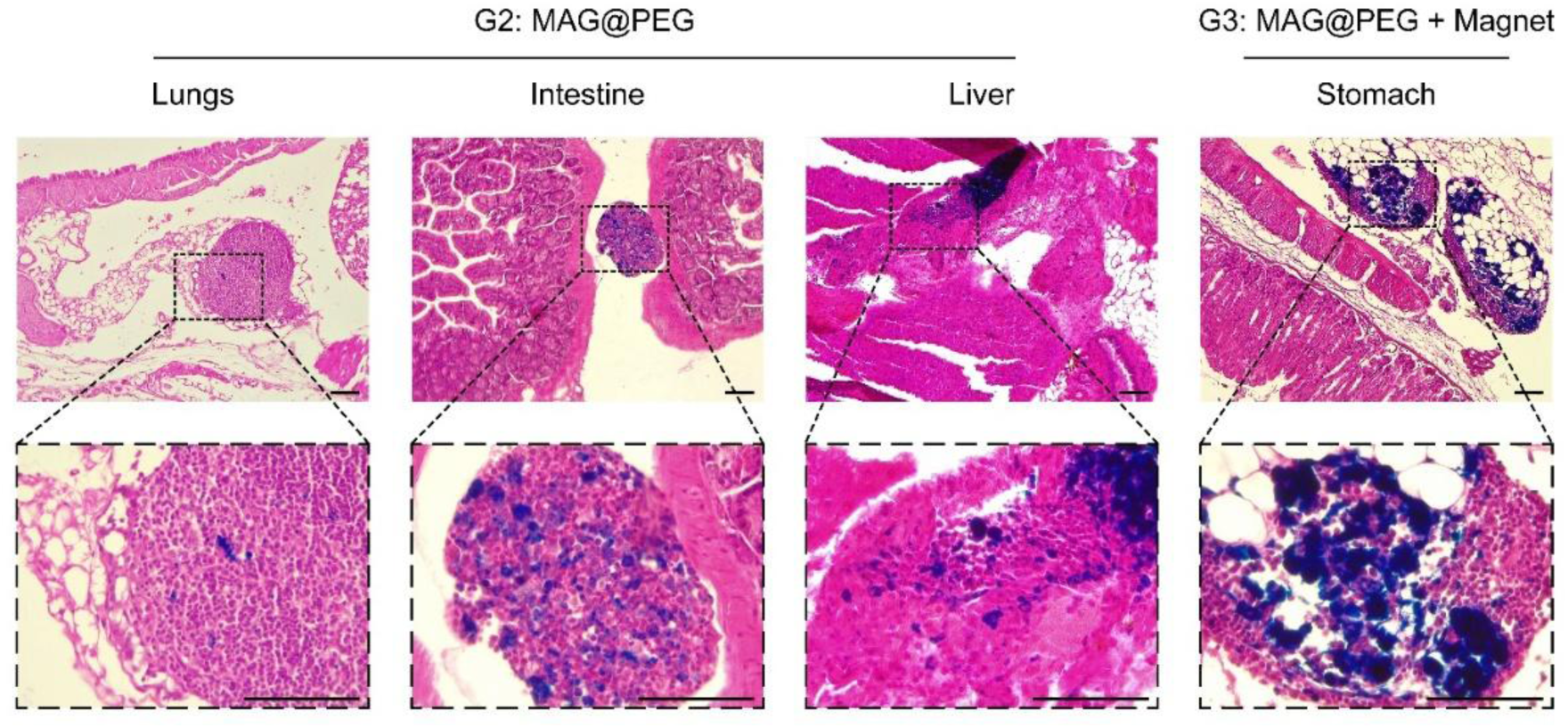
Presence of Prussian blue-positive deposits in organ-associated lymphoid follicles. Prussian blue positive deposits of varying sizes were observed in lymphoid follicles associated with the lungs, intestine, and liver (G2), and lymphoid-adipose structures in the stomach (G3). Scale bar: 100 µm.

### 3.3. Hemocompatibility of MAG@PEG in human blood

Following the *in vivo* evaluation, the hemocompatibility of MAG@PEG was investigated using human blood samples to further assess their biosafety for potential clinical application (**Figure 7**).

**Figure 7.**
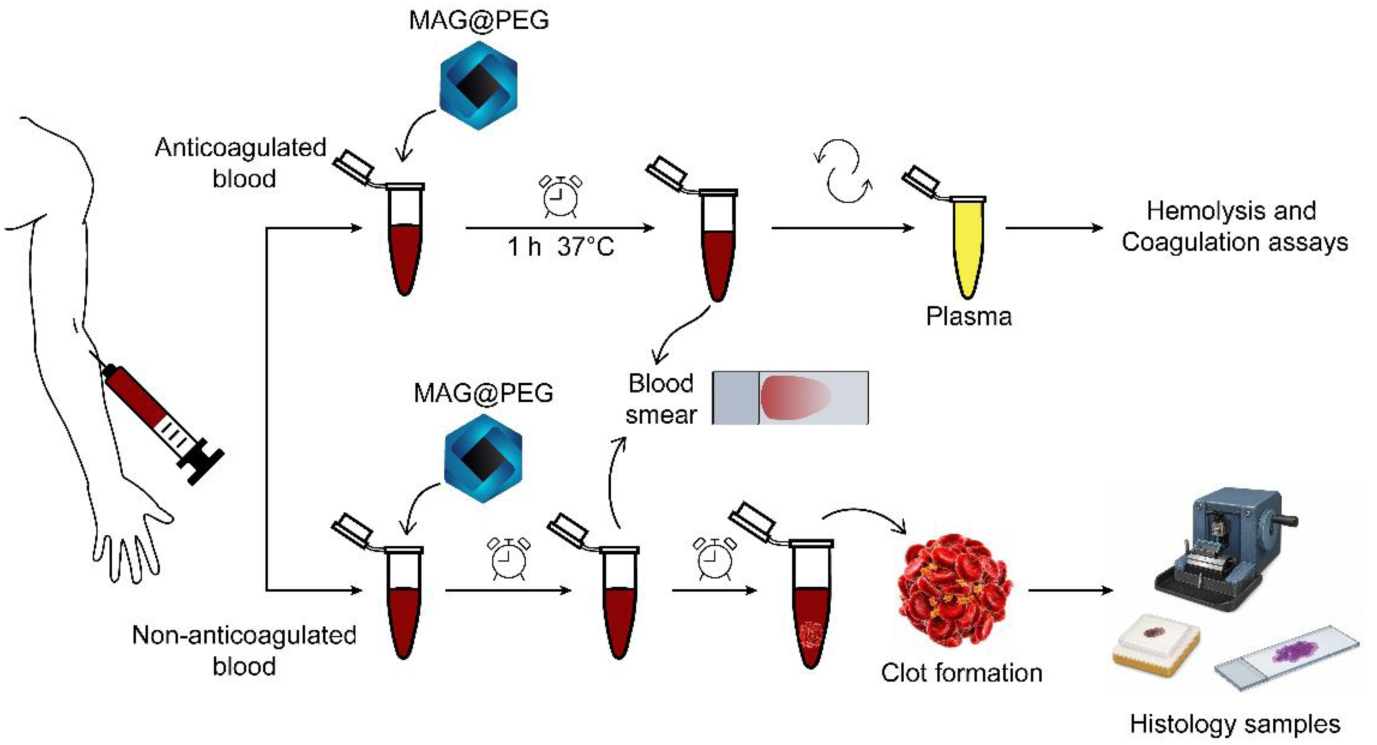
Experimental design for hemocompatibility assessment in human blood. Anticoagulated whole blood was incubated with MAG@PEG at 37 °C for 1 h. Plasma was then separated to perform hemolysis and coagulation assays. In parallel, non-anticoagulated blood samples were used to assess clot formation and to prepare blood smears for morphological and histological analysis.

Firstly, peripheral blood smears were prepared from anticoagulated human blood samples, and cell morphology was observed by optical microscopy. Red blood cells incubated with vehicle (control) or 160 µg/mL of MAG@PEG displayed normal size, color, and morphology (**Figure 8A**).

**Figure 8.**
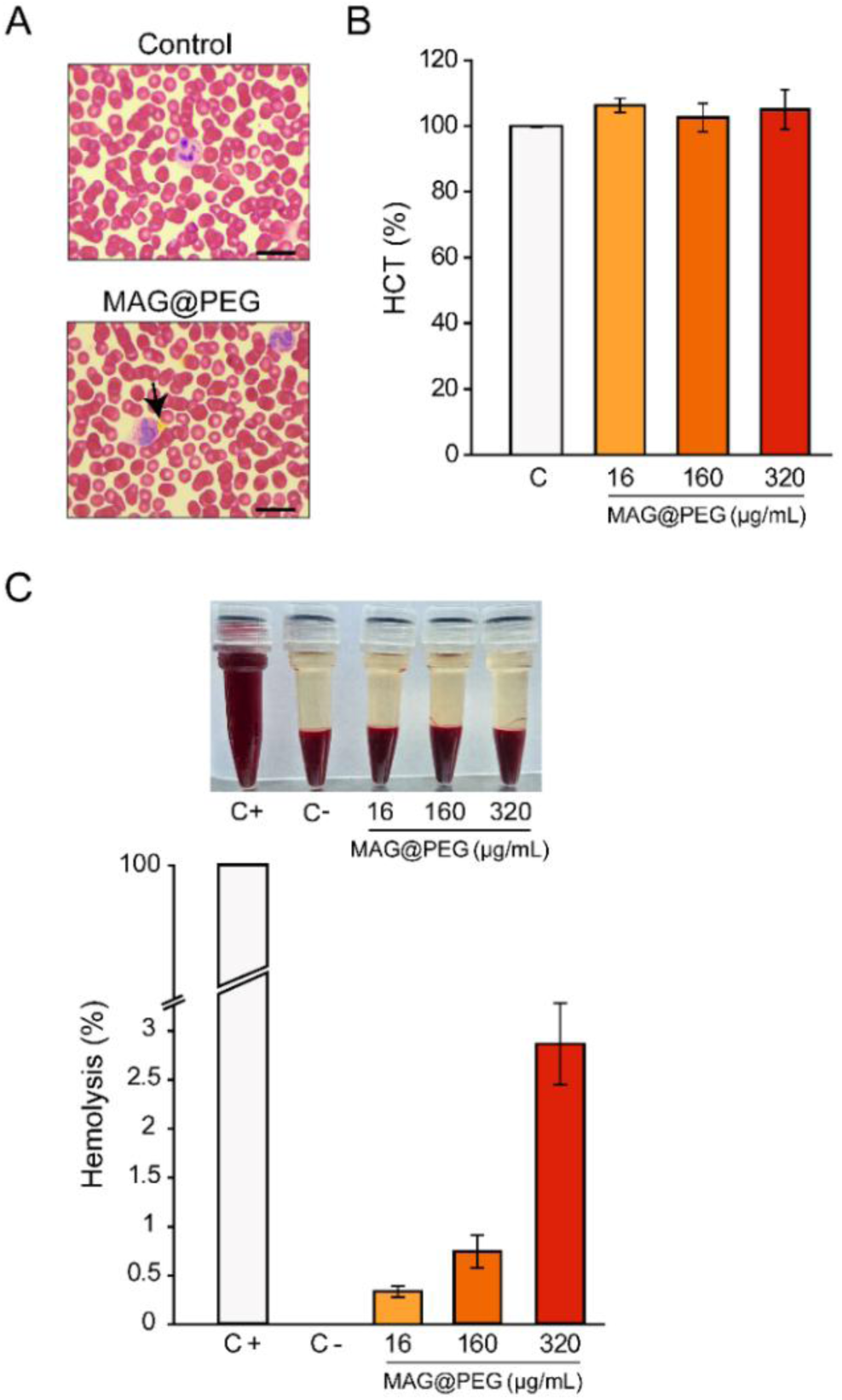
**Hemolysis assay**. Peripheral blood smears (A) and hematocrit (B) were prepared from anticoagulated human blood after 1 h incubation at 37 °C with physiological saline (control) or 160 µg/mL MAG@PEG. Representative images of smears are shown (black arrow: MAG@PEG; scale bar: 20 µm). (C) Anticoagulated human blood was incubated with increasing concentrations of MAG@PEG (16, 160, and 320 µg/mL) for 1 h at 37 °C, and hemoglobin release was measured spectrophotometrically at 540 nm. Results are expressed as percentage of control (mean ± SD, n = 5). Statistical analysis was performed using a linear mixed-effects model followed by Tukey’s test (*p < 0.05; **p < 0.01; ***p < 0.001).

Furthermore, neither red blood cell distribution nor morphological features were affected in the presence of MAG@PEG (black arrow). White blood cells and platelets also exhibited normal morphology under both conditions. Then, hematocrit values determined after incubation with increasing MAG@PEG concentrations (16-320 µg/mL) showed no statistically significant differences compared to the control, indicating that MNP exposure does not affect the erythrocyte volume fraction under the tested conditions (**Figure 8B**).

Secondly, the *in vitro* hemocompatibility of MAG@PEG was evaluated following the ASTM standard protocol for assessing the hemolytic properties of materials (ASTM F 756-00). According to this standard, materials are classified as non-hemolytic (< 2% hemolysis), slightly hemolytic (2-5%), or hemolytic (> 5%). As shown in **Figure 8C**, blood samples in the presence of MAG@PEG exhibited low hemolysis levels, with no significant differences among conditions. Although the highest concentration tested (320 µg/mL) could be classified as slightly hemolytic, all values remained comparable to the negative control, supporting an overall non-hemolytic profile.

Furthermore, the effect of MAG@PEG on the activation of the coagulation cascade and clot formation was evaluated to complement the hemocompatibility assessment. Thrombin time (TT), activated partial thromboplastin time (aPTT), prothrombin time (PT), and fibrinogen levels in plasma samples were incubated with increasing concentrations of MAG@PEG (16-320 µg/mL). As shown in **Figure 9A**, TT, PT, and fibrinogen values remained unchanged; while aPTT was significantly prolonged at the highest concentration. These indicate that MAG@PEG at concentrations below to 160 µg/mL, do not significantly interfere with the intrinsic or extrinsic coagulation pathways. Nevertheless, aPTT values at 320 µg/mL of MAG@PEG suggests that elevated levels of iron may exert some influence on the intrinsic and/or common coagulation pathway.

**Figure 9.**
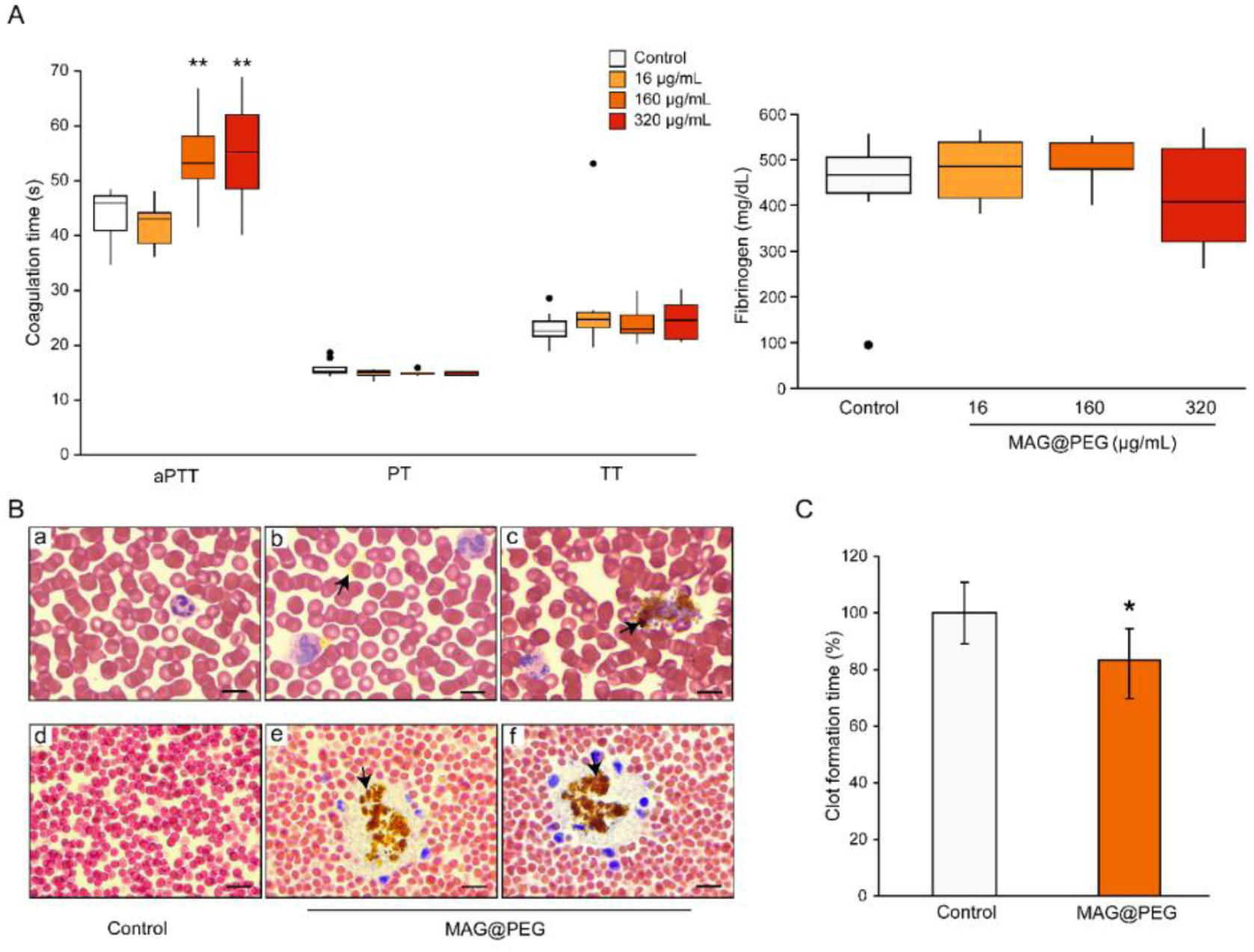
Coagulation assays. (A) Plasma samples were incubated with increasing concentrations of MAG@PEG (16-320 µg/mL), and the coagulation parameters thrombin time (TT), activated partial thromboplastin time (aPTT), prothrombin time (PT) and fibrinogen were assessed. (B) Whole blood analysis of clot formation. Upper panel: representative peripheral blood smears showing anticoagulated blood in the absence (a; control) or presence of 160 µg/mL MAG@PEG (b), and non-anticoagulated blood exposed to 160 µg/mL MAG@PEG (c). Lower panel: histological sections of clots showing control clot (d) and clots formed in the presence of 160 µg/mL MAG@PEG (e and f). (C) Clot formation time was quantified after exposure to 160 µg/mL of MAG@PEG. Results are expressed as percentage of control (mean ± SD, n = 5). Statistical analysis was performed using a linear mixed-effects model followed by Tukey’s test for (A) (*p < 0.05; **p < 0.01) and Student’s t-test for (C) (*p < 0.05).

In addition, platelet-MNPs interactions were found to be dependent on anticoagulant presence. In anticoagulated blood both, platelets and MNPs, were homogeneously distributed throughout the smear after 1 h of incubation with MAG@PEG (**Figure 9B, b**). In contrast, in non-anticoagulated blood, the preincubation with MAG@PEG led to the formation of platelet-MNP aggregates (**Figure 9B, c**). In both conditions, the morphology and distribution of red and white blood cells remained comparable to those observed in the control samples.

Representative histological sections of a clot formed in the absence or presence of 160 µg/mL MAG@PEG are exhibited in **Figure 9B, d-f**. The control sample showed normal clot architecture, characterized by a fibrin network with evenly distributed blood cells (**Figure 9B, d**). In contrast, clots formed in the presence of MAG@PEG displayed a fibrin network associated with white blood cells and platelets surrounded by aggregates of MNPs, suggesting interactions between these blood components and the nanosystem (**Figure 9B, e** and **f**). Notably, red blood cells did not appear to be involved in this effect. Finally, clot formation time in non-anticoagulated blood exposed to 160 µg/mL of MAG@PEG was found significantly decreased compared to the control (**Figure 9C**).

## 4. Discussion

The physicochemical properties of MAG@PEG support their suitability for *in vivo* applications. In particular, their hydrodynamic diameter, size distribution, and surface charge indicate good colloidal stability, while also being key determinants of biodistribution and circulation behavior, as reported both in our previous studies and by other authors [22, 23, 24].

In the present study, we evaluated the biodistribution and safety of the MAG@PEG nanosystem in tumor-bearing mice, with and without the application of an external magnet to promote tumor accumulation. As mentioned above, the administered dose of MAG@PEG (10 mg/kg) was selected based on a previous report by Agotegaray and col. (2017). In that study, the administration of chitosan-coated magnetic nanoparticles at a dose of 30 mg/kg produced signs of potential renal damage [21], as histopathological examination of kidney sections revealed abnormal areas compatible with periarteriolar nephritis. In contrast, in the present investigation, no histopathological or biochemical alterations indicative of renal damage were observed at the selected dose of 10 mg/kg (Table 1 and Fig. 5). Moreover, this study evaluated the biodistribution of MAG@PEG after one month of treatment and one week after the final administration (Fig. 1). Under these conditions, Prussian blue-positive staining was not detected in most organs, except for the spleen and in lymphoid-associated structures in the lung, stomach and intestine (Fig. 5 and 6). In this regard, the spleen consistently exhibited a higher Prussian blue area in the treated groups (G2 and G3) compared with the control group (Suppl. Fig. 3). The presence of Prussian blue-positive staining in spleen sections from the control group may be explained by the ability of the spleen to filter the blood and remove senescent erythrocytes, leading to the accumulation of iron-containing degradation products, particularly in the red pulp [25]. Furthermore, this physiological function of the spleen may also explain MAG@PEG retention, which was consistent with the second-highest accumulation detected by VSM (Fig. 3B). This finding indicates that at least a fraction of the nanosystem reached the systemic circulation following intraperitoneal administration. At the same time, visible amounts of MAG@PEG were observed during mice dissection (Suppl. Fig. 4), suggesting that a fraction of the administered MNPs remained within the peritoneal cavity. Notably, this retention was observed in both G2 and G3 mice, making it unclear whether the application of the magnet influenced MAG@PEG distribution and/or accumulation.

There is evidence indicating that changes in the colloidal stability of the nanosystem may produce MNPs agglomeration and local retention [26]. This phenomenon could partially explain the higher magnetization values detected in organs from G2 when compared with G3 (Fig. 3A), as well as the greater inhibitory effect on tumor development observed in G2 (Fig. 2C). In this regard, the antitumor effect detected in G2 may be associated with a systemic outcome of the MAG@PEG administration rather than with MNP accumulation in the tumor area induced by magnetic targeting. On the other hand, the presence of Prussian blue-positive deposits in tumor sections from G3 (Fig. 4) may be related to the immediate application of the magnet following the intraperitoneal injection. Under these conditions, the magnetic field could have influenced the initial distribution of MAG@PEG, promoting MNP accumulation in the tumor area while favoring partial retention within the peritoneal cavity. This interpretation is consistent with the visible MNP retention observed during dissection (**Suppl. Fig. 5**) and the lower magnetization values detected in several organs from G3 compared with G2 (Fig. 3A). Similar effects associated with intraperitoneal administration have been reported when compared with intravenous administration [27, 28].

As reported in other investigations, the liver and the spleen are usually the major sites of MNPs accumulation [29, 30]. However, their biodistribution depends mostly on the size [31] and the surface characteristics of the nanosystem [26, 32], which influence their recognition by macrophages of the mononuclear phagocytic system (MPS) [33]. In contrast, the stomach was the organ that displayed the highest mass fraction of MAG@PEG in the present study (Fig. 3). This finding may be attributed either to a local effect of the intraperitoneal route of administration [34] or to the accumulation of MNPs in lymphoid-adipose tissue (Fig. 6), which is enriched in macrophages [35]. Additionally, smaller Prussian blue-positive deposits detected in lymphoid follicles associated with the lungs, intestine, and liver (Fig. 6) may be related to the presence of Kupffer cells in the liver [36] and other macrophages of the MPS in the intestine and lungs [28]. In any case, the presence of these iron deposits did not appear to substantially alter the normal tissue architecture when compared with the control group. In relation to this observation, the nanosystem did not produce evident adverse effects in mice, as overall behavior, body weight (Fig. 2A), and biochemical parameters (Table 1) remained largely unchanged. Although a statistically significant increase in liver transaminases was detected in G3, which may suggest potential hepatocellular damage, no differences were observed in either the histological analysis (Fig. 5) or in the magnetization measurements (Fig. 3A).

Several medical applications require direct contact of MNPs with blood and healthy tissues. It has been extensively discussed in the literature that poorly designed nanostructures can induce hemolysis or cytotoxicity due to protein binding, cellular interaction, and phagocytosis [16, 37]. One strategy to improve the biological performance of MNPs is their functionalization with non-ionic surfactants and biocompatible polymers. In this regard, PEG has been widely employed because of its ability to improve biocompatibility and reduce nanoparticle clearance *in vivo* [11]. In the present study, we further evaluated the hemocompatibility of MAG@PEG for its potential application as a multifunctional nanoplatform. The results showed that exposure of human blood to MAG@PEG did not affect leukocyte, platelet, or erythrocyte morphology. Furthermore, no significant changes in hematocrit values were observed with increasing concentrations of MAG@PEG (Fig. 8). Moreover, MAG@PEG was classified as non-hemolytic according to the ASTM F756 standard. In agreement with this finding, Tween 80-PEG 3350-coated MNPs have also been reported to be safe in biological systems at low concentrations [37, 38]. Thus, PEG-coating may prevent direct contact of the metallic core with the erythrocyte membrane, drastically reducing toxic interaction and hemolysis.

Bare MNPs typically accelerate coagulation by shortening PT and/or aPTT due to their charged surface that activates Factor XII, whereas PEG-coating and other polymeric coatings can prevent these effects [37, 39]. Consistent with these reports, MAG@PEG did not markedly alter the conventional coagulation parameters evaluated in this study. Although a moderate prolongation of aPTT was observed at the highest concentrations tested, PT, TT, and fibrinogen levels remained unaffected (Fig. 9A) Furthermore, blood smear and clot histology analyses did not reveal evident morphological alterations associated with MAG@PEG exposure (Fig. 9B). These findings suggest that PEG-coating may reduce the interaction between MAG@PEG with plasma proteins and cellular blood components, contributing to its hemocompatibility.

## 5. Conclusion

Taken together, the combined biodistribution, histological, biochemical, and hemocompatibility findings provide a comprehensive preclinical assessment of the biosafety of MAG@PEG and identify the route of administration and magnetic targeting as key determinants of nanoparticle biodistribution.

## Acknowledgments

This work was supported by grants awarded to V.G-P. from Consejo Nacional de Investigaciones Científicas y Técnicas (CONICET PIP 11220210100811CO), Universidad Nacional del Sur (PGI 24/B305 and 24/B364), and Agencia Nacional de Promoción Científica, Tecnológica y de la Innovación (PICT-2020-1922), Argentina.

## Notes

### Competing Interest Statement

The authors have declared no competing interest.

