## Supplementary Material for "Preclinical evaluation of the biodistribution, safety, and hemocompatibility of PEG-coated iron oxide magnetic nanoparticles in a viral oncogenesis mouse model"

[a]Depto. de Biología, Bioquímica y Farmacia, Universidad Nacional del Sur (UNS), San Juan 670, 8000, Bahía Blanca, Argentina.

[b]Instituto de Ciencias Biológicas y Biomédicas del Sur (INBIOSUR), UNS-Consejo Nacional de Investigaciones Científicas y Técnicas (CONICET), San Juan 671, 8000, Bahía Blanca, Argentina.

[c]Depto. de Química, Universidad Nacional del Sur (UNS), Av. Alem 1253, 8000, Bahía Blanca, Argentina.

[d]Instituto de Química del Sur (INQUISUR), UNS-CONICET, Av. Alem 1253, 8000, Bahía Blanca, Argentina.

[e]Instituto de Física La Plata (IFLP), UNLP-CONICET, Diagonal 113 entre 63 y 64, 1900, La Plata, Argentina.

[f]Bioquímica Austral, Laboratorio de Análisis Clínicos y Gestión, 25 de Mayo 1007, 8000, Bahía Blanca, Argentina.

[#]These authors contributed equally

*Corresponding author: Verónica González-Pardo,

Departamento de Biología, Bioquímica y Farmacia, UNS.

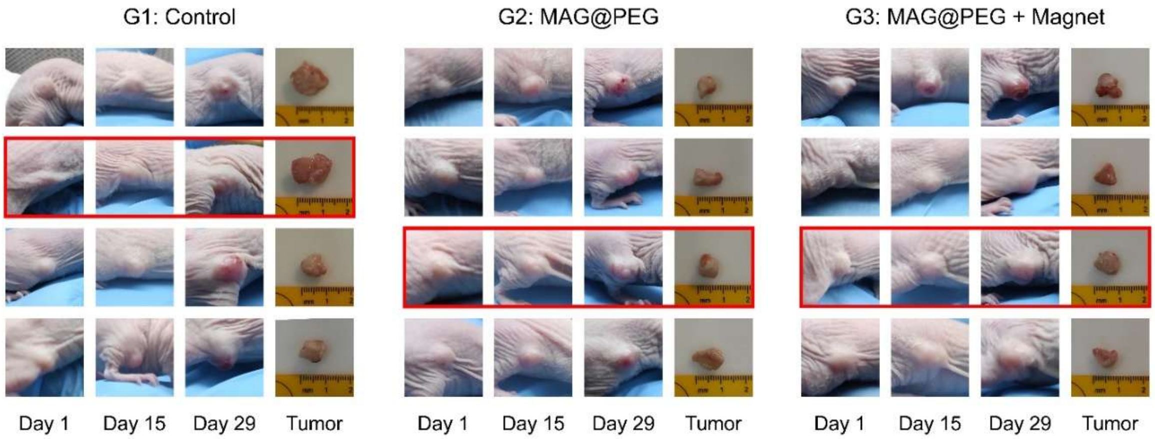


**Suppl. Fig. 1. Representative images of tumor progression throughout the experimental period.** Representative photographs of mice on treatment days 1, 15, and 29 and their corresponding excised tumors are shown. Red rectangles indicate the image sets for presentation in Figure 1.

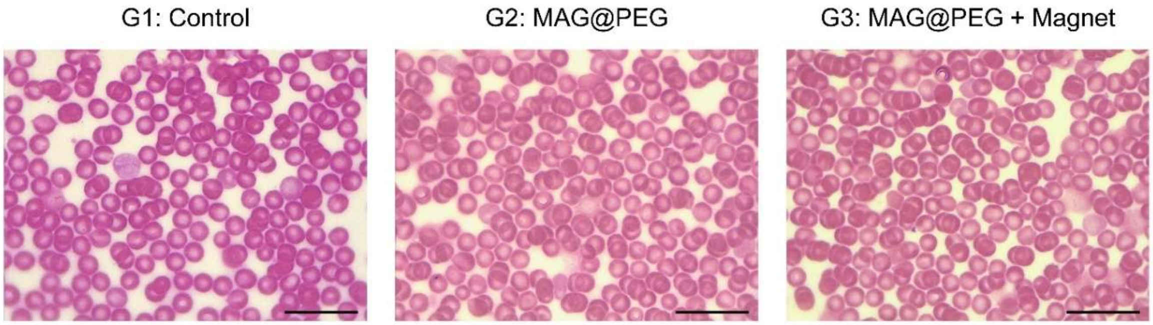


**Suppl. Fig. 2. Representative blood smears from treated mice.** Representative peripheral blood smears obtained from mice in the Control (G1), MAG@PEG (G2), and MAG@PEG + Magnet (G3) groups at the end of the experiment. Scale bar = 20 μm.

### Determination of biodistribution and accumulation of MNPs in tumors and organs

#### MNP mass fraction from Saturation magnetization

Two main features of the M vs. H curves are the quasi-linear regions observed in the high- and low-field regions, hereafter referred to as the high- and low-field susceptibilities, respectively (Supp. Fig. 3.I).

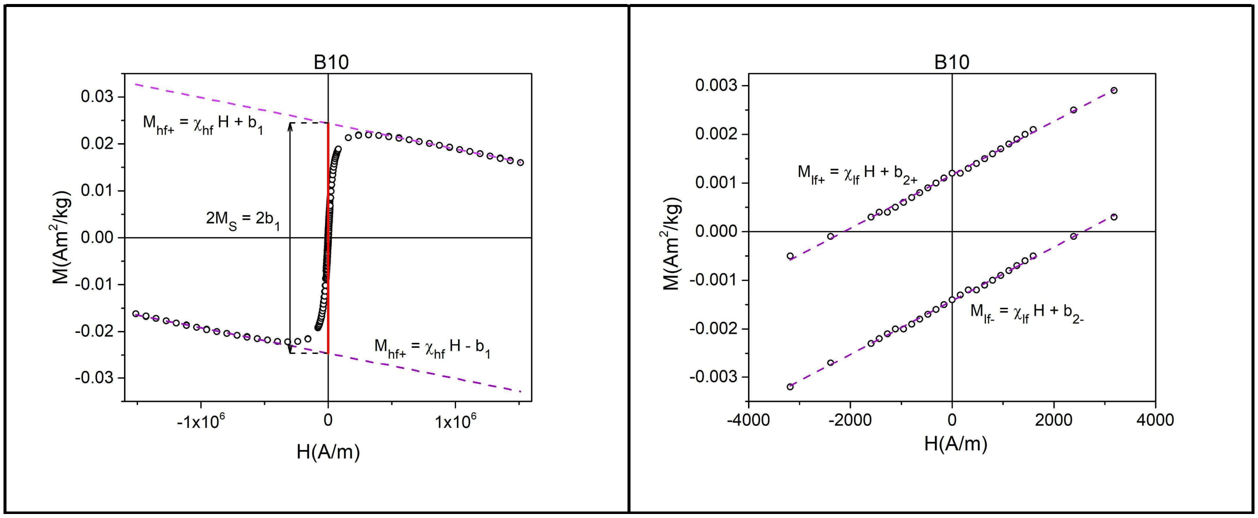


**Suppl. Fig. 3.I. Quasi - linear regions of the magnetization curve.**

The high-field (hf) magnetizations on both branches are well approximated by

$M_{hf+} = \chi_{hf} H + b_1$

$M_{hf-} = \chi_{hf} H - b_1$

where $\chi_{hf}$ is the high field susceptibility. The parameter $b_1$ approaches well the saturation magnetization $M_S$, i.e., $M_S \approx b_1$

From the sample and MNP powder, the mass fraction of MNP in the sample, $x = m_{MNP} / m_S$, can be estimated, where $m_{MNP}$ and $m_S$ are the masses of MNP and tissue in the sample, respectively.

In effect, $M_S = \mu_S / m_S = \mu_S(MNP) / m_S$, where $\mu_S$ stands for the moment of the MNP in the sample. Using $x = m_{MNP} / m_S$,

$m_S = m(MNP) / x$,

and

$M_S = x^* \mu_S(MNP) / m(MNP) = x^* M_S(MNP) \Rightarrow x = M_S / M_S(MNP)$

#### Mass fraction from low field susceptibility

The low-field (lf) susceptibility are the slopes of the magnetization curves in the quasi-linear low field region,

$M_{lf+} = \chi_{lf} H + b_{2+}$

$M_{lf-} = \chi_{lf} H + b_{2-}$

Neglecting the minor effect due to the approach to saturation, the high-field susceptibility is essentially the diamagnetic susceptibility of the tissues, $\chi_{hf} \sim \chi_{dia}$. The low-field susceptibility is the sum of the contributions from the MNPs, $\chi$, and from the tissues, $\chi_{dia}$, such that $\chi_{lf} = \chi + \chi_{dia}$. Hence,

$$\chi = \chi_{lf} - \chi_{hf}$$

The value of $\chi$ is proportional to the mass concentration of MNPs, x', such that

$$x' = \chi / \chi(MNP)$$

where $\chi$(MNP) is the value recorded for the MNP powder. The following figure shows the correlation between the x' values and the $x = M_S / M_S(MNP)$ values used in eq. (1) of the main manuscript.

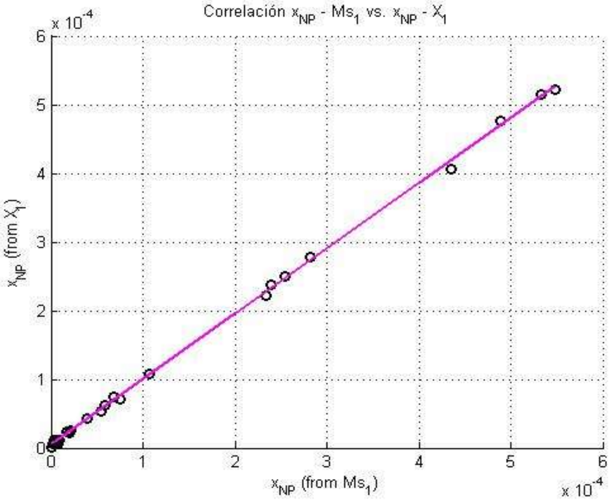


**Suppl. Fig. 3.II. Correlation between mass fractions obtained from Ms and from $\chi$**

The excellent correlation between the mass fractions estimated from Ms and χ, together with the slope of the linear fit (0.95), which is close to unity, provides strong evidence that the MAG MNPs retained their magnetic properties after *in vivo* administration.

**Comparison of coercivity in tissues and in MNP powder**

The high-field magnetization ($M_{hf}$) as a function of the field ($H_{hf}$) on the positive (+) and negative (-) branches was approximated as

$$M_{hf+} = \chi_{hf} H + b_1$$

$$M_{hf-} = \chi_{hf} H - b_1$$

For the low-field magnetization, it has been approximated

$$M_{lf+} = \chi_{lf} H + b_{2+}$$

$$M_{lf-} = \chi_{lf} H + b_{2-}$$

where $b_{2\pm}$ are the intercepts of both lines.

The magnetization due exclusively to the MNPs is

$M(MNP)_{lf+} = \chi H + b_{2+}$

$M(MNP)_{lf-} = \chi H + b_{2-}$

It follows that

$M(MNP)_{lf+} = M_{lf+} - \chi_{hf} * H$

The condition for determining the coercive field (H = Hc) is $M(MNP)_{lf+} = 0$, i.e.,

$M_{lf+} - \chi_{hf} Hc = 0,$

that is,

$\chi_{lf} Hc + b_{2+} - \chi_{hf} Hc = (\chi_{lf} - \chi_{hf}) Hc + b_{2+} = 0,$

hence,

$Hc_{+} = -b_{2+} / (\chi_{lf} - \chi_{hf})$

Similarly,

$Hc_{-} = -b_{2-} / (\chi_{lf} - \chi_{hf})$

In the following table, the coercivities obtained using this procedure, from tissues displaying strong enough signal, and from MNP powder, are listed.

| Tissue | Hc (A/m) | SD (A/m) |
|---|---|---|
| Spleen | 2342 | 12 |
| Stomach | 2769 | 6 |
| Lung | 2389 | 35 |
| MAG MNPs | 2664 | 8 |

**Suppl. Table 1. Average coercivities in tissues (groups 2 and 3) and MAG powder**

From these results, a mean value for the tissues was obtained:

$Hc_m = \sum Hc_i/SD_i / \sum 1/SD_i = 2595$ A/m

where the statistical weights are the inverses of the SDs. The error associated to $Hc_m$ is:

$SD_m = (\sum((Hc_m - HC_i)^2/SD_i) / \sum 1/SD_i)^{½} = 206$ A/m

The percentage difference between this average and the coercivity of the MNP powder (2664 A/m) is less than 3%. Suppl. Fig. III shows the distribution of these results together with the fitted normal distribution.

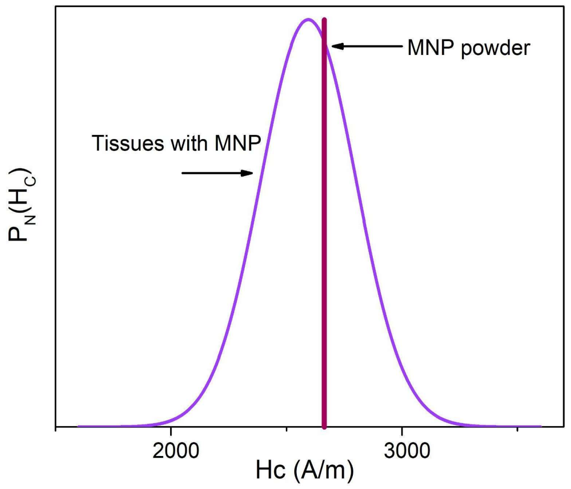


**Suppl. Fig. 3.III. Coercivities in tissues and MNP powder**

Together, the analyses presented in this section confirm that the MAG MNPs retain their magnetic properties, supporting the reliable estimation of their mass fraction in tissues using Eq. (1) of the main manuscript.

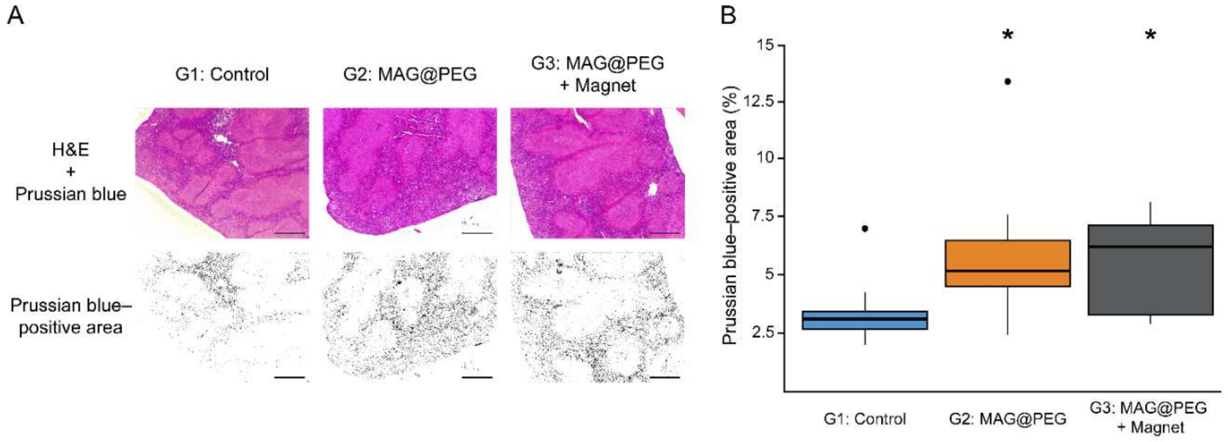


**Suppl. Fig. 4. Quantification of Prussian blue-positive staining in the spleen.** (A) Upper panel: Representative spleen micrographs stained with H&E and Prussian blue. Lower panel: Prussian blue-positive areas were defined using the Image J program. (B) The bar graph represents the quantification of the Prussian blue-positive area across the different groups. Significant differences were evaluated using one-way ANOVA followed by Dunnett's post hoc test (*$p < 0.05$). Scale bar: 500 µm.

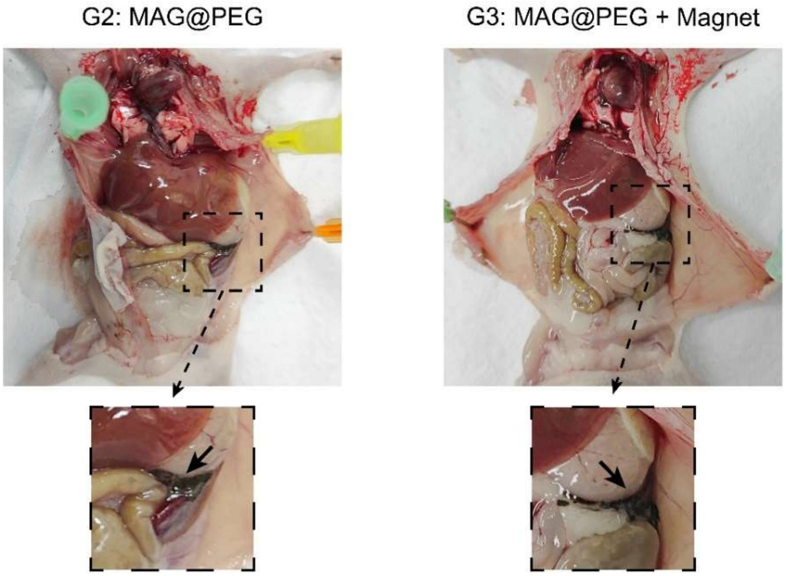


**Suppl. Fig. 5. Macroscopic evidence of MNP retention in the peritoneal cavity.** Representative photographs obtained during dissection of MAG@PEG (G2) and MAG@PEG + magnet (G3) mice are shown. Dashed rectangles indicate enlarged areas highlighting visible nanoparticles retention within the peritoneal cavity.
